# SpaPheno: Linking Spatial Transcriptomics to Clinical Phenotypes with Interpretable Machine Learning

**DOI:** 10.1101/2025.09.18.676993

**Authors:** Bin Duan, Xiaojie Cheng, Hua Zhou

## Abstract

Linking spatial transcriptomic data to clinically relevant phenotypes is essential for advancing spatially informed precision oncology. Here, we present SpaPheno, an interpretable machine learning framework that integrates spatial transcriptomics with clinically annotated bulk RNA-seq to identify spatially resolved biomarkers predictive of patient outcomes, including survival, tumor stage, and immunotherapy response. SpaPheno provides multi-scale interpretability from tissue regions to cell types and individual spatial spots, enabling clear biological insights from complex spatial data. We validate SpaPheno through extensive simulations and applications to multiple cancer cohorts—primary liver cancer, clear cell renal cell carcinoma, breast cancer, and melanoma—demonstrating robust predictive performance alongside biologically meaningful spatial patterns. SpaPheno offers a generalizable strategy to translate spatial omics data into clinically actionable knowledge, facilitating precision oncology informed by tumor spatial architecture.

## Introduction

Recent advances in spatial transcriptomics (ST) have revolutionized our ability to resolve gene expression landscapes within intact tissues, offering unprecedented insights into cell–cell interactions, tissue architecture, and disease pathology at near-cellular resolution[1]. These technologies are increasingly applied across biomedical domains—from developmental biology to cancer research and regenerative medicine[2]. Yet, a critical challenge remains unresolved: how can spatial transcriptomic data be systematically leveraged to inform clinically relevant decisions?[3, 4] [5]

Translating spatial molecular patterns into patient-level clinical insights faces three fundamental challenges. First, clinical phenotypes such as survival, tumor stage, or therapy response are typically associated with bulk RNA-seq cohorts that lack spatial resolution, making it non-trivial to infer spatial correlates of clinical traits. Second, ST data are often sparse, noisy, and platform-dependent, complicating robust modeling across samples and regions[6]. Third, clinical application demands not only predictive performance but also biological interpretability—particularly for identifying actionable targets or guiding spatially informed interventions[7, 8]. Recent efforts have started to explore the linkage between spatial omics and clinical phenotypes. For example, SpaLinker[9] decomposes bulk transcriptomic profiles into latent factors using non-negative matrix factorization (NMF), associates these factors with clinical outcomes, and subsequently maps their distributions back to spatial transcriptomic data. In contrast, stClinic[10] employs a dynamic graph model with Gaussian mixture priors to delineate spatial niches and leverages attention-based learning to connect niche-level representations with clinical features in datasets where spatial and clinical information are paired. While these approaches represent important advances, they also have notable limitations: SpaLinker essentially performs a global bulk-based factorization and only visualizes the inferred factors in spatial data without explicitly modeling spatial organization, whereas stClinic relies on rare cohorts with matched clinical and spatial information, limiting its generalizability across cancer types and larger clinical cohorts.

To address these challenges, we developed SpaPheno, an interpretable machine learning framework designed to uncover spatially localized regions and cell types predictive of clinical outcomes. SpaPheno bridges bulk RNA-seq data annotated with clinical labels and spatial transcriptomic data lacking phenotypes, enabling spatial biomarker discovery through three key innovations. First, SpaPheno constructs biologically interpretable, low-dimensional features by integrating cell-type composition with local spatial context, allowing both spatial transcriptomics and bulk RNA-seq data to be embedded into a shared, cell-type–resolved feature space. Second, it employs Elastic Net regression(Enet)—a sparsity-aware, interpretable model well-suited to high-dimensional data with correlated features[11]. Third, it integrates SHapley Additive exPlanations (SHAP)[12] to assign interpretable importance scores to spatial and cellular features, enabling localized biological insights. Importantly, SpaPheno is not a simple sequential combination of Elastic Net and SHAP; rather, it constitutes an integrated framework in which feature construction, sparse modeling, and interpretable attribution are jointly designed to yield biologically meaningful, multi-scale insights from spatial omics data.

We benchmarked SpaPheno using two cortex spatial transcriptomic datasets[13, 14] with simulated phenotypes and real-world datasets from four tumor types—primary liver cancer[15, 16], clear cell renal cell carcinoma (ccRCC)[17], breast cancer (BRCA)[18], and melanoma[19]—with clinical phenotypes including overall survival, tumor stage, and immune checkpoint blockade (ICB) response. Across all settings, SpaPheno consistently demonstrates high predictive accuracy and strong biological interpretability. Together, these results establish SpaPheno as a generalizable and interpretable framework for linking spatial molecular features to clinical phenotypes, supporting spatially informed precision oncology.

## Results

### 1. Overview of the SpaPheno

SpaPheno is an interpretable machine learning framework that links spatial transcriptomic features to clinical phenotypes by integrating spatial and bulk RNA-seq data (**Fig. 1**). It addresses three major challenges in translational spatial modeling: (i) the lack of spatial–clinical paired datasets, (ii) the sparsity and noise of spatial transcriptomic measurements, and (iii) the need for biologically interpretable models to support clinical understanding. To overcome these challenges, SpaPheno operates in three stages: (1) unified feature representation of spatial and bulk data, (2) phenotype prediction and spatial projection, and (3) multi-scale interpretation via SHAP-based analysis.

**Fig. 1.**
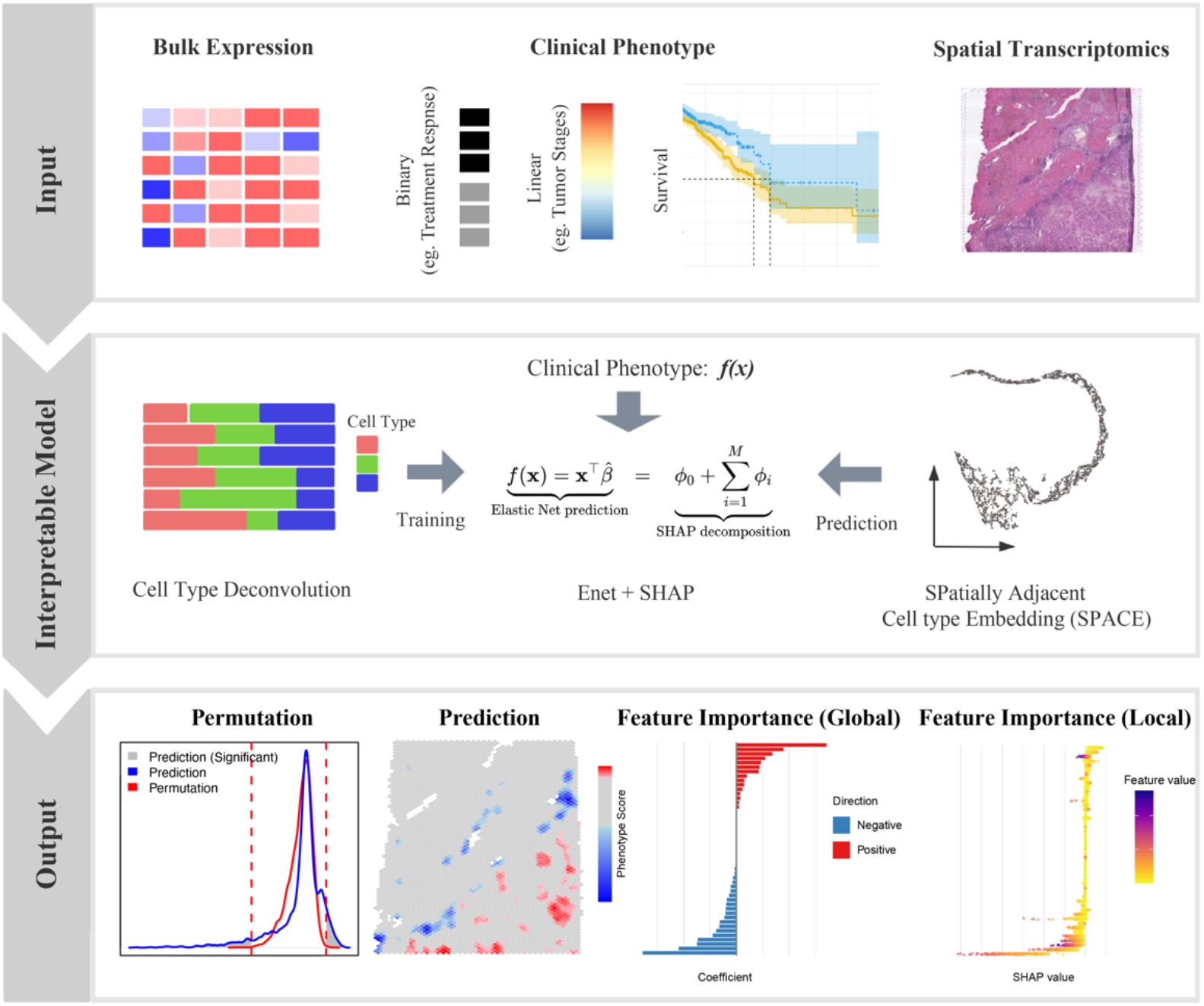
Workflow of SpaPheno. SpaPheno consists of three modules: **Input**, which includes bulk RNA-seq data with clinical phenotypes and spatial transcriptomics (ST) data; **Interpretable model**, which integrates the two data modalities via a shared cell type–based feature space, derives spatial embeddings (SPACE), and applies an Enet model with SHAP for interpretable phenotype prediction; **Output**, which highlights spatial regions associated with clinical phenotypes and provides both global and local feature attributions for biological interpretation.

First, SpaPheno takes as input (i) bulk RNA-seq cohort with clinical phenotypes and (ii) spatial transcriptomic profiles. Both datasets are projected into a shared cell-type– based feature space using a common single-cell reference, enabling consistent and biologically interpretable representations across data modalities[20, 21]. To incorporate spatial context, after obtaining cell-type-based feature of ST data, we apply our previously developed SpaDo[6] algorithm to derive spatially adjacent cell-type embeddings (SPACE), which summarize local cell-type neighborhoods for each spot. These embeddings enhance robustness by smoothing local variability while preserving tissue microenvironmental heterogeneity—critical for detecting localized, clinically relevant niches.

Using the unified cell-type–based representations, SpaPheno trains an Elastic Net (Enet) regression model on the bulk RNA-seq data to associate spatial patterns with clinical phenotypes such as overall survival, tumor stage, or immunotherapy response. We adopted Enet here, because it balances feature selection and regularization, especially under correlated inputs. The trained model is then applied to spatial data to generate spot-level predictions of clinical relevance. To assess the specificity of spatial predictions, we implement a permutation-based significance test by randomly shuffling spatial coordinates and recalculating prediction scores, helping to distinguish biological signal from spatial autocorrelation artifacts (see Methods).

Finally, SpaPheno computes SHAP values at the spot level to reveal spatial drivers of clinical phenotypes. These values quantify the contribution of each cell type within each spatial unit to the predicted clinical outcome, enabling multi-scale interpretability— from tissue regions, to cell types, to individual spots. By focusing on regions with significantly high or low SHAP scores, SpaPheno identifies spatial outliers that may correspond to immune niches, fibrotic zones, or other pathologically relevant microenvironments. This final step transforms SpaPheno from a predictive model into a powerful tool for spatial biomarker discovery and biological interpretation.

### 2. Simulation on real single cell spatial transcriptomics

To systematically evaluate SpaPheno, we conducted simulation studies grounded in real spatial transcriptomic datasets, primarily osmFISH[13]and STARmap[14]. They provide both cell-type annotations and spatial domain labels, enabling realistic phenotype simulations and quantitative benchmarking. Specifically, we simulated binary phenotypes by randomly selecting pairs of anatomically defined layers and aggregating cell-type compositions to mimic bulk RNA-seq profiles (**Fig. 2a-b, see Methods**). To better reflect real-world scenarios in which phenotype classes are not perfectly separated, we included comparisons between spatial domains with subtle distinctions—such as Lay3-median versus Lay3-lateral—which belong to the same layer and are difficult to distinguish. Biological heterogeneity was further introduced by incorporating background signals from other spatial regions and applying mild stochastic perturbations to the cell-type proportions. Each simulation yielded 50 synthetic bulk samples per class, capturing inter-sample variability at the cohort level (see **Methods**). SpaPheno was then trained on these simulated bulk samples and evaluated for its ability to recover spatially localized phenotype differences.

**Fig. 2.**
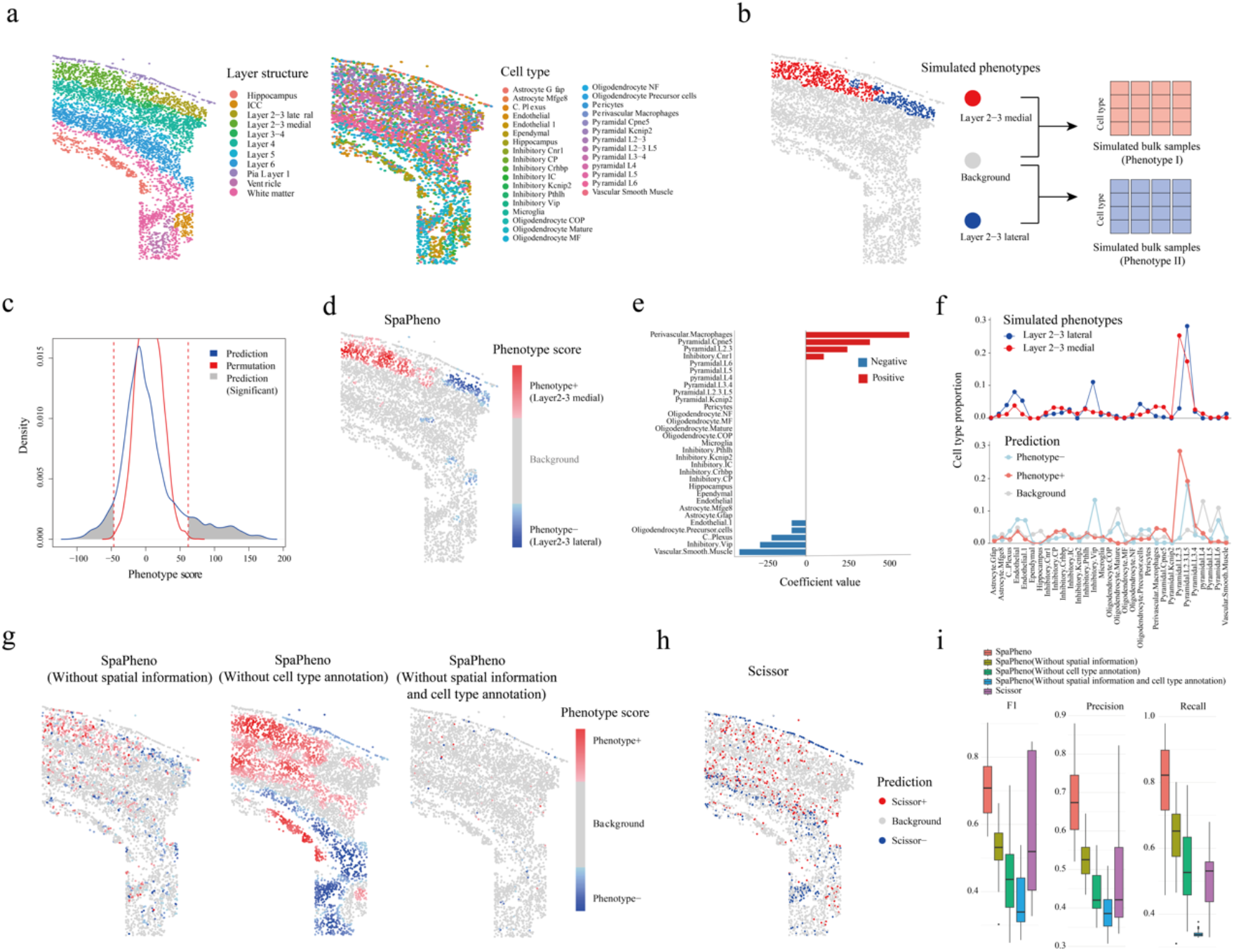
Evaluation of SpaPheno on simulated phenotypes using real osmFISH data. **a**, Layer structure and cell type annotations in the osmFISH dataset. **b**, Simulation of bulk RNA-seq samples and binary phenotype labels. **c**, Distribution of SpaPheno-predicted phenotype scores compared with permutation controls. **d**, Spatial prediction map of phenotype scores across the tissue. **e**, Global feature attributions based on model coefficients. **f**, Comparison of cell type composition between simulated and predicted phenotype groups. **g**, Performance under ablation settings removing spatial information, cell-type information, or both. **h**, Prediction results from Scissor. **i**, Overall performance comparison of SpaPheno, its ablation variants, and Scissor across all simulated phenotype scenarios.

Our results show that SpaPheno successfully identified phenotype-associated spatial regions with significant enrichment (p < 0.001, permutation test by shuffling cell coordinates), and the predicted regions aligned well with ground truth annotations (**Fig. 2b-d**). The model’s coefficient values and the cell-type compositions of predicted phenotype regions were also highly consistent with the simulated signals (**Fig. 2e–f**), indicating that SpaPheno can effectively capture phenotype-relevant differences in cell-type composition and precisely map them onto spatial coordinates.

One advantage of SpaPheno is to design unified representation by combining cell-type annotation and spatial information. To assess the value of the two components (cell-type annotation and spatial information) of SpaPheno, we implemented three ablation models: (i) using only cell-type annotation without spatial information; (ii) using spatially smoothed gene expression without cell-type decomposition; (iii) a naive model with neither spatial nor cell-type features. In addition, we further benchmarked against Scissor[22], a state-of-the-art phenotype linkage method developed for single-cell data. In a particularly challenging comparison between the Lay3-median and Lay3-lateral, SpaPheno consistently outperformed all ablations and Scissor, demonstrating its effectivity and superiority (**Fig. 2g-h**).

To test the robustness of SpaPheno, we extended the simulation framework across all pairwise combinations of layers in the osmFISH dataset (**Fig. 2i**). Additionally, we assessed the impact of spatial smoothing by varying the number of neighbors used in the SPACE embedding (**Supplementary Fig. 1**). To further evaluate generalizability, we replicated the simulations and ablations using the STARmap[14] dataset, confirming its broad applicability across spatial transcriptomic technologies (**Supplementary Fig. 2**).

In addition to its predictive power, SpaPheno offers multi-scale interpretability—from global feature attributions to region-level insights, and down to individual cell types and spatial spots (**Fig. 2e, Supplementary Fig. 3, and see Methods**). This multi-scale interpretability enhances model transparency and supports biological interpretation.

### 3. SpaPheno identifies survival-associated spatial regions across multiple tumor types

To demonstrate SpaPheno’s applicability in real-world clinical contexts, we applied it to spatial transcriptomic datasets from three cancer types—primary liver cancer[15, 16], breast cancer (BRCA)[18], and clear cell renal cell carcinoma (ccRCC)[17]—selected to represent increasing tumor microenvironmental complexity. We first focused on patient survival as the primary phenotype due to its broad clinical relevance.

In primary liver cancer, which includes four hepatocellular carcinoma (HCC) slices, one combined HCC-cholangiocarcinoma (cHCC) slice, and one intrahepatic cholangiocarcinoma (ICC) slice, SpaPheno consistently identified low-risk survival-associated regions enriched in tertiary lymphoid structures (TLS), known for their dense B and T cell aggregates and favorable prognosis[23] (**Fig. 3, Supplementary Fig. 4-5**). Using SHAP analysis, we quantified the contributions of individual cell types to survival-associated regions, where negative SHAP values indicated association with low-risk survival and positive values indicated high-risk. SpaPheno revealed distinct survival-related cellular patterns, exemplified by cytotoxic CD8 T cells enriched in low-risk regions and SPP1^+^ macrophages prevalent in high-risk regions, consistent with prior studies[16, 24] (**Fig. 3b-c, Supplementary Fig. 5a**). Notably, cell-type contributions to low-risk survival regions were highly consistent across spatial slices, suggesting a conserved immune microenvironment associated with better outcomes (**Fig. 3b**). In contrast, high-risk survival regions exhibited greater heterogeneity in both cell composition and SHAP attribution patterns, reflecting diverse, context-dependent tumor-promoting microenvironments (**Fig. 3c**). This contrast underscores SpaPheno’s ability to resolve not only predictive regions, but also their biological coherence or divergence across spatial contexts.

**Fig. 3.**
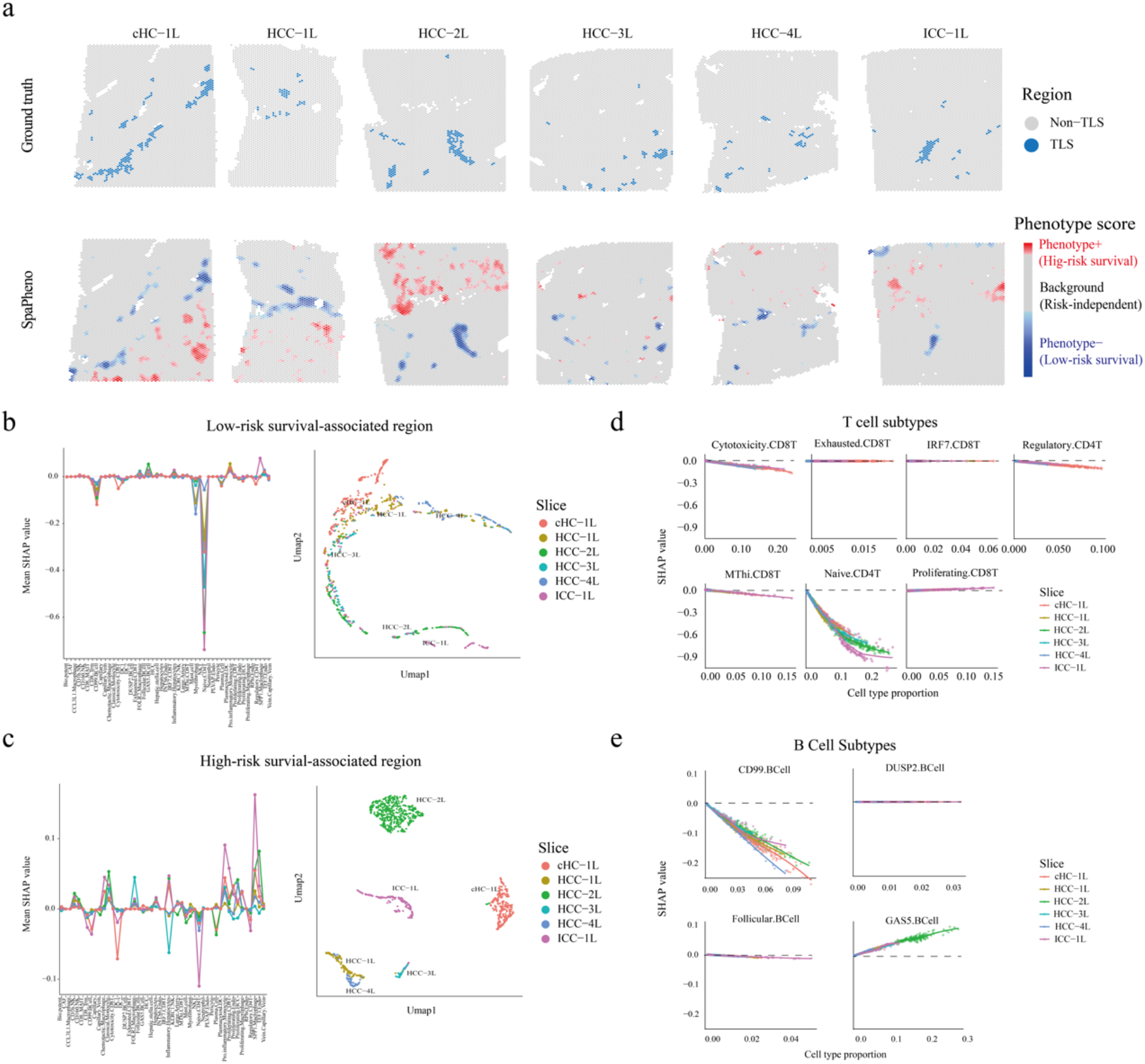
SpaPheno links TLS-like niches to low-risk survival in primary liver cancer. **a**, TLS annotations and SpaPheno-predicted survival-associated regions across six ST slices. **b**, Mean SHAP values of each cell type and UMAP visualization of predicted low-risk spots by their SHAP value of each cell type. **c**, Mean SHAP values and UMAP visualization of high-risk spot by their SHAP value of each cell type. **d**, SHAP values of T cell subtypes in low-risk survival-associated regions across slices; lower SHAP values indicate lower predicted risk. **e**, SHAP values of B cell subtypes in low-risk survival-associated regions across slices.

In addition, SpaPheno uncovered functional heterogeneity within major immune cell types. For instance, not all T cells were associated with favorable prognosis—only naive CD4 T cells exhibited clear protective effects, while other subtypes, such as exhausted CD8 T cells showed no association (**Fig. 3d, Supplementary Fig. 5c**). Similarly, among B cells, CD99+ B cells were linked to better survival, whereas GAS5+ B cells correlated with poor outcomes (**Fig. 3e, Supplementary Fig. 5c**). These patterns were not limited to B and T cells—other lineages also displayed subtype-specific effects (**Supplementary Fig. 5b-c**)—emphasizing SpaPheno’s capacity to dissect fine-grained phenotypic associations beyond canonical cell classes. These findings are consistent with previous studies demonstrating the prognostic relevance of immune cell functional heterogeneity in tumors[25-27].

To further validate these insights, we applied SpaPheno to an independent HCC dataset[16]. Here, high-risk survival-associated regions predicted by SpaPheno coincided with areas enriched in SPP1+ macrophages, while low-risk regions corresponded to immune-rich niches (**Supplementary Fig. 6**), underscoring the robustness and generalizability of the model.

Extending beyond liver cancer, we tested SpaPheno in BRCA[18] and ccRCC[17], which represent increasing tumor microenvironmental complexity. For BRCA, SpaPheno revealed survival-associated spatial patterns similar to those observed in HCC, with low-risk regions co-localizing with immune infiltration zones (**Supplementary Fig. 7**). For ccRCC, SpaPheno identified two distinct classes of low-risk survival regions—one enriched for TLS-like immune structures and another characterized by endothelial cell enrichment but relative lower T and B cells (**Supplementary Fig. 8**), suggesting a potential role of vascularization in contributing to favorable prognosis in ccRCC. Together, these results highlight SpaPheno’s ability to uncover survival-associated spatial regions across diverse cancer types.

### 4. Spatial correlates of tumor stage reflect immune architecture in primary liver cancer

To further demonstrate SpaPheno’s utility in characterizing clinical phenotypes, we analyzed its performance in identifying tumor stage–associated regions across primary liver cancer samples (**Fig. 4, Supplementary Figs. 9–11**). Similar to the survival analysis, SpaPheno was trained on bulk RNA-seq data labeled by early versus late tumor stage, then spatial patterns were inferred across multiple liver cancer slices.

**Fig. 4.**
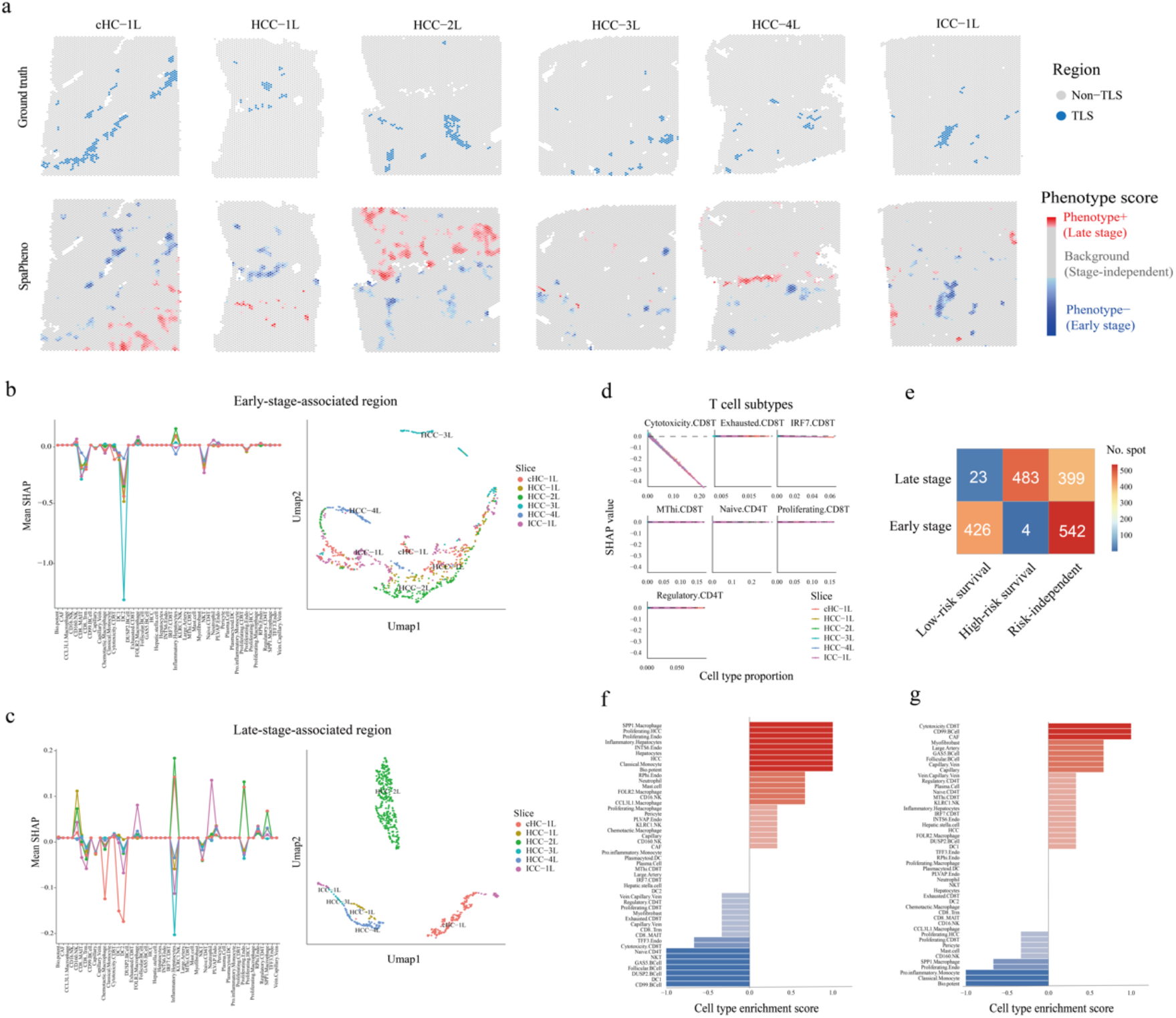
SpaPheno distinguishes tumor stage–associated TLS-like regions from survival-independent niches. **a**, TLS annotations and SpaPheno-predicted stage-associated regions across six ST slices. **b**, Mean SHAP values of each cell type and UMAP visualization of early-stage– associated regions. **c**, Mean SHAP values and UMAP visualization of late-stage–associated regions. **d**, SHAP values of T cell subtypes in early-stage–associated regions across slices; lower SHAP values indicate lower predicted stage. **e**, Confusion matrix comparing survival- and stage-associated spatial regions. **f**, Cell type composition of early-stage–specific, survial-independent regions. **g**, Cell type composition of late-stage–specific, survival-independent regions.

Consistent with known biology, regions associated with early tumor stage frequently overlapped with TLS–like areas, characterized by dense immune infiltration (**Fig. 4a, d, Supplementary Fig. 10**). To systematically quantify cell-type contributions, SHAP analysis was performed on six representative slices. Negative SHAP values indicated association with early stage, while positive values indicated late stage. We found that cell-type contributions linked to early-stage regions were largely consistent across patients (**Fig. 4b**), whereas late-stage regions exhibited greater heterogeneity in cellular composition (**Fig. 4c**). These results suggest that tumor progression is accompanied by increasing spatial and cellular complexity.

Notably, spatial predictions associated with tumor stage and survival showed strong concordance: early-stage-associated regions commonly overlapped with low-risk survival-associated zones, whereas late-stage-associated regions aligned with high-risk survial-associated areas, further validating SpaPheno’s ability to capture clinically relevant features (**Fig. 4e**). However, two distinct discordant region types emerged: (i) early-stage–specific but survival-independent regions, and (ii) late-stage–specific but survival-independent regions. To elucidate these, we examined representative examples. Early-stage–specific, survival-independent regions were enriched for proliferating HCC cells and SPP1^+^ macrophages, but exhibited limited immune infiltration (**Fig. 4f, Supplementary Fig. 11a**), suggesting that despite early pathological staging, such regions may harbor elevated risk and warrant closer monitoring. Conversely, late-stage–specific, survival-independent regions showed strong infiltration of cytotoxic CD8^+^ T cells and depletion of SPP1^+^ macrophages (**Fig. 4g; Supplementary Fig. 11b**), indicating active anti-tumor immunity and potentially less aggressive clinical behavior than expected.

Together, these findings highlight SpaPheno’s capacity to provide spatially resolved, nuanced insights beyond conventional tumor staging, with potential implications for personalized treatment strategies.

### 5. SpaPheno identifies ICB-responsive immune niches in melanoma

We next evaluated SpaPheno’s ability to identify spatial determinants of immune checkpoint blockade (ICB) response using a published melanoma spatial transcriptomics dataset[19] integrated with bulk-defined responder and non-responder labels[28]. As this dataset lacked predefined spatial domains, we first applied BayesSpace[29] to infer spatial clusters and then incorporated histopathological annotations from H&E images to assign region identities at the spot level (**Fig. 5a–c**).

**Fig. 5.**
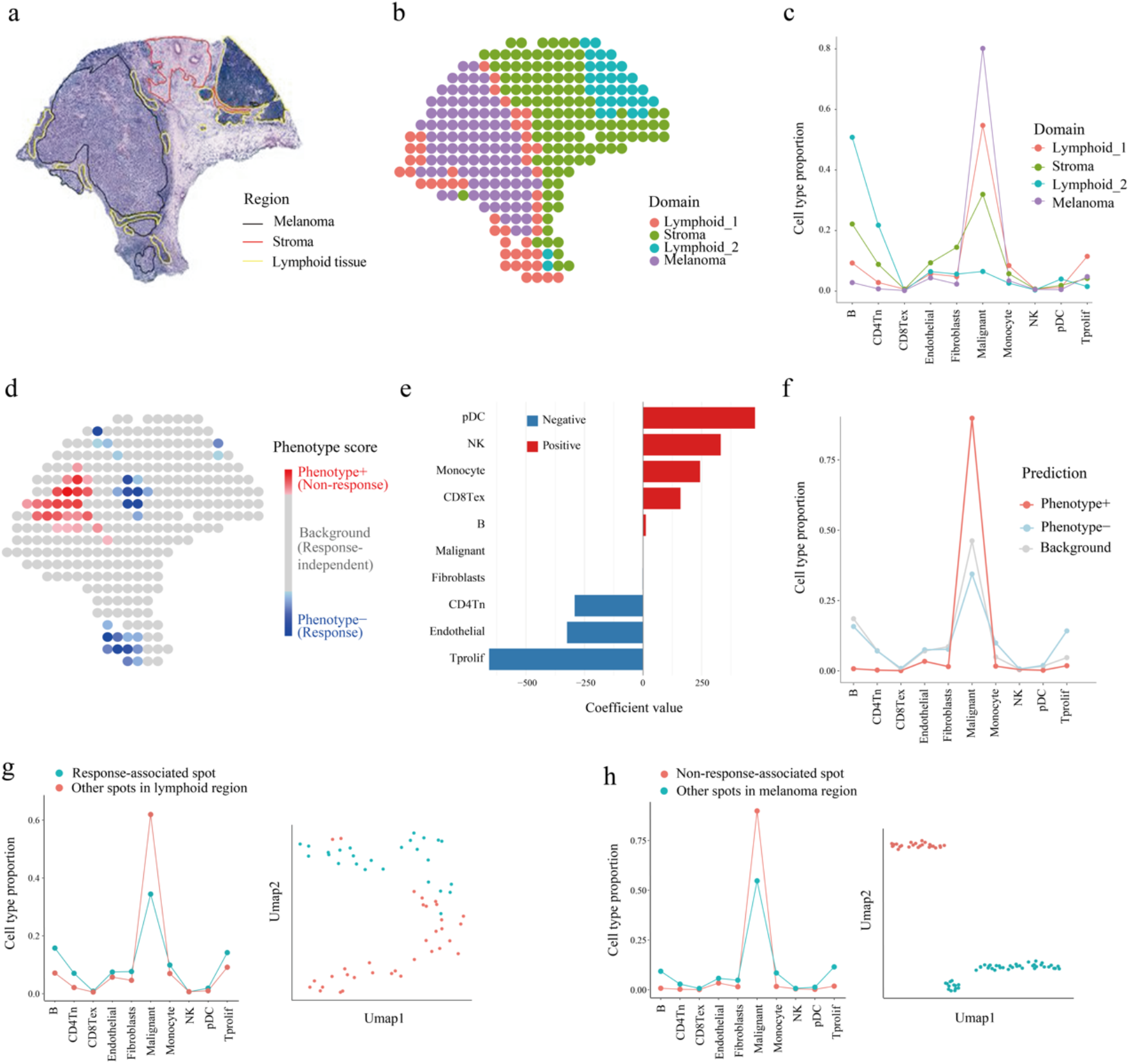
SpaPheno maps ICB response–associated immune niches in melanoma. **a**, H&E staining and region annotations of a melanoma tissue slice. **b**, Spatial domain annotations for each spot. **c**, Cell type composition across domains. **d**, SpaPheno-predicted spatial distribution of ICB response scores. **e**, Model coefficients indicating global cell type contributions to ICB response. **f**, Cell type composition of SpaPheno-predicted response versus non-response regions. **g**, Cell type composition and UMAP visualization of ICB response–associated versus other spots within lymphoid regions. **h**, Cell type composition and UMAP visualization of ICB non-response–associated versus other spots within melanoma regions.

SpaPheno revealed that specific immune cell types, such as proliferative T cells (Tprof), were positively associated with treatment response, whereas exhausted CD8^+^ T cells (CD8^+^ Tex) correlated with non-response (**Fig. 5e–f**). Importantly, SpaPheno identified the “Lymphoid_1” domain as the primary immune-enriched region associated with ICB response, while the other “Lymphoid_2” domain, despite also being immune-rich, exhibited little or no association with treatment response (**Fig. 5d**).

Further analysis of these two domains revealed key differences in cellular composition and spatial localization. “Lymphoid_1” was notably enriched with both B cells and T cells and was spatially situated at the tumor boundary. In contrast, “Lymphoid_2” contained markedly fewer B and T cells and was located farther away from the tumor region. This suggests that effective ICB response requires a spatial niche characterized by both high infiltration of multiple immune cell types—especially T and B cells—and close proximity to tumor cells.

Moreover, non-responder regions localized to tumor areas with reduced immune infiltration despite similar tumor cell density, underscoring the importance of immune microenvironment composition and spatial context in shaping therapeutic outcomes.

Together, these findings demonstrate SpaPheno’s ability to dissect immune subtype-specific and spatially resolved features that underlie ICB response. By moving beyond bulk immune content, SpaPheno provides mechanistic insights into spatial immune niches critical for immunotherapy efficacy, highlighting its utility in spatial immuno-oncology research.

## Discussion

In this study, we present SpaPheno, an interpretable machine learning framework that bridges spatial transcriptomic data with clinically annotated bulk RNA-seq profiles. By integrating Elastic Net regression with SHAP-based feature attribution, SpaPheno enables spatial localization and mechanistic interpretation of clinically relevant signals, including patient survival, tumor stage, and immunotherapy response. Applied across multiple cancer types—liver, breast, kidney cancers, and melanoma—SpaPheno consistently identified phenotype-associated spatial regions and revealed their underlying cellular contributors.

Unlike black-box predictive models or purely descriptive spatial analyses, SpaPheno offers multi-scale interpretability. It not only achieves accurate phenotype prediction but also explains where and why these associations arise, through region-, cell-type-, and spot-level SHAP contributions. The framework’s robust performance across diverse tumor types and clinical endpoints highlights its translational potential.

Another key advantage of SpaPheno is its sensitivity to subtle, spatially confined biological differences. For example, it can detect clinically discordant regions—areas within a tumor that differ in prognosis- or treatment-relevant features—thereby offering new opportunities for refining diagnosis and improving risk stratification. By disentangling phenotype-related spatial signatures in a biologically grounded and explainable manner, SpaPheno facilitates deeper insights into the cellular architecture and functional heterogeneity of the tumor microenvironment.

Importantly, while demonstrated here primarily in cancer spatial transcriptomics, SpaPheno’s design is broadly applicable to other diseases and biological contexts where spatial transcriptomic data and corresponding bulk phenotype information are available. This generalizability positions SpaPheno as a versatile tool for deciphering spatial phenotype associations beyond oncology, potentially extending to neurological disorders, inflammatory diseases, and developmental biology.

Looking forward, SpaPheno lays the groundwork for several promising directions. These include extending the framework to multi-sample or longitudinal spatial datasets, integrating other data modalities such as spatial proteomics or single-cell multi-omics, and applying it in prospective clinical cohorts for biomarker discovery and patient stratification. As spatial technologies move closer to clinical translation, interpretable frameworks like SpaPheno will be essential for ensuring that spatially informed predictions can guide decision-making in precision oncology and beyond.

## Methods

### Interpretable phenotype-associated spatial signature identification by SpaPheno

The workflow of SpaPheno is illustrated in **Fig. 1**. SpaPheno integrates two types of input data: (1) a bulk RNA-seq gene expression matrix with clinical phenotype vector, (2) a spatial transcriptomics matrix. The phenotype can be continuous (e.g., survival time), binary (e.g., treatment response), or categorical (e.g., tumor stage).

To enable phenotype association analysis in a comparable feature space, SpaPheno first apply cell2location[21] to deconvolve both the bulk and spatial transcriptomics data using the same reference single-cell RNA-seq dataset. For the spatial transcriptomics data, we further enhance the feature representation by incorporating spatial context. Specifically, SpaPheno adopts our previously developed SPACE (Spatially Adjacent Cell-type Embedding) framework to compute spatial neighborhood-aware cell type distributions for each spot. This results in a spatially enriched cell-type feature matrix where each spot is represented by the average of cell-type proportions in its spatial neighborhood. This embedding reflects both local composition and tissue structure, and effectively transforms each spot into a spatially-aware pseudo-bulk sample, improving comparability with the bulk profiles.

With both bulk and spatial data now represented in a shared cell-type feature space, SpaPheno proceeds to fit a phenotype-predictive regression model using Enet regularization (Fig. 1). We use linear regression for continuous outcomes, logistic regression for binary outcomes, and cox proportional hazards regression for survival data. The learned model is then used to predict phenotype scores for each spatial spot, highlighting regions most associated with the phenotype of interest.

Finally, SpaPheno applies SHAP to interpret the model’s predictions, quantifying the contribution and directionality of each cell type to the phenotype score at both global and local levels. Additional residual-based analyses identify biologically significant regions such as potential hotspots or conserved zones.

### Elastic Net modeling and phenotype score calculation

To establish a quantitative link between cell-type compositions and clinical phenotypes, SpaPheno fits an elastic net regression model using deconvoluted bulk RNA-seq data as the input feature matrix and a clinical phenotype vector as the response. The elastic net combines the strengths of both L1 (lasso) and L2 (ridge) penalties, providing sparse yet stable solutions particularly suitable for high-dimensional compositional data.

Let

- *X*^*bulk*^∈ ℝ^*n*×*p*^ be the cell-type proportion matrix for *n* bulk samples across *p* cell types;
- *Y* ∈ ℝ^*n*^ be the phenotype vector, which can be continuous, binary, or survival-based.

The elastic net objective is formulated as:

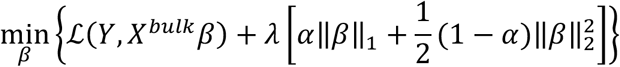

where

- ℒ is the loss function (e.g., squared loss for continuous phenotypes, logistic loss for binary classification, or partial likelihood for Cox regression),
- *λ* controls the overall penalty strength, and
- α ∈ [0,1] balances sparsity and stability.

After fitting the model on bulk data, the learned coefficient vector 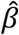 is used to calculate phenotype scores for each spatial spot *i*, based on its corresponding SPACE feature vector 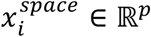.

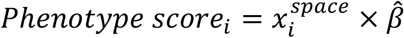

These phenotype scores reflect the degree to which each spatial region exhibits the phenotype of interest. Importantly, this prediction step is model-driven and does not involve SHAP, which is used purely for post hoc interpretation.

Specifically, we applied Elastic Net (Enet) regression using the cv.glmnet function from the R package glmnet to identify features associated with clinical phenotypes. The Enet model linearly combines L1 (lasso) and L2 (ridge) regularization, with a mixing parameter alpha controlling their relative contributions. For survival phenotypes, we set family = “cox”; for binary outcomes (e.g., ICB response), we used family = “binomial”; and for continuous or categorical phenotypes (eg. tumor stage), we used family = “gaussian”.

To automatically select the optimal values of *α* and *λ*, we predefined a grid of candidate alpha values ranging from 0.005 to 0.9 (*α* ∈ {0.005,0.01,0.05,0.1, …,0.9}). For each alpha, 10-fold cross-validation was performed using function cv.glmnet of R package glmnet, and the mean cross-validation error (cvm) at lambda.min (i.e., the value of lambda that minimizes the error) was recorded. The alpha corresponding to the lowest cvm was selected as the optimal mixing parameter. A final Enet model was then trained using this optimal alpha and its corresponding lambda.min. All procedures were performed with a fixed random seed (seed = 2025) to ensure reproducibility.

### SHAP for interpretable spatial contribution scores

To interpret the contributions of different cell types to the predicted phenotype scores, SpaPheno applies SHAP, a unified framework grounded in cooperative game theory. SHAP decomposes each model output into additive feature contributions, enabling both global (feature importance) and local (spot-level reasoning) explanations. Specifically, all SHAP and interpretability analyses were performed using the R package iml.

For each spatial spot *i*, SHAP calculates the prediction *ŷ*_*i*_ as:

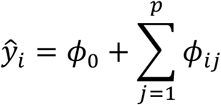

Where *ϕ*_0_ is the global bias term and *ϕ*_*ij*_ represents the contribution of cell type *j* to the prediction at spot *i*.

SpaPheno employs several SHAP-based visualization strategies:

- Summary plot: Provides an overview of the distribution and direction of cell-type contributions across the entire tissue. This helps identify which cell types are consistently associated (positively or negatively) with the phenotype across space.
- Dependency plot: Plots SHAP value against the abundance of a specific celltype, revealing nonlinear or interaction-driven effects. This enables detailed exploration of how the impact of a cell type changes with its local abundance.
- Waterfall plot: By inspecting SHAP values at individual spots, SpaPheno provides a local decomposition of phenotype scores, enabling fine-grained reasoning at the spot level.

### Residual-based interpretation of SHAP values

To move beyond direct feature attribution, SpaPheno incorporates residual analysis on SHAP dependence plots to uncover phenotype-associated spatial outliers. Specifically, for each cell type, we fit a linear model between its abundance and the corresponding SHAP values:

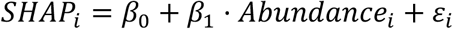

where *ε*_*i*_ denotes the residual for spot *i*. We then compute the standardized residuals (z-scores) across all spots:

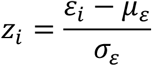

Spots with *z*_*i*_ > 2 or *z*_*i*_ < −2 are identified as SHAP outliers:

- High-positive residuals (*z*_*i*_ > 2) indicate biological hotspots, where the phenotype contribution is higher than expected given cell type abundance.
- High-negative residuals (*z*_*i*_ < −2) suggest conserved zones, where the phenotype impact is lower than expected.

These residual outliers reveal nonlinear effects or context-specific modulations of cell type–phenotype relationships, offering novel insights into spatial heterogeneity. By bridging predictive modeling with interpretable residual patterns, SpaPheno enables biologically grounded hypothesis generation and facilitates downstream experimental validation.

### Permutation-based significance analysis

To evaluate whether the spatial phenotype associations identified by SpaPheno reflect true spatial organization rather than random chance, we perform a permutation-based significance test. This involves randomly shuffling the spatial coordinates of all spots to disrupt spatial structure, followed by recalculating the SPACE embeddings and predicting phenotype scores using the elastic net model trained on bulk data. This generates a null distribution of predicted scores representing spatially random scenarios.

Significant phenotype-associated spots are defined as those with predicted scores exceeding empirical thresholds *p* derived from this null distribution. By default, we use a two-sided threshold corresponding to the top and bottom *p* of the null distribution (By default, *p* = 0.001).

Because the threshold is based on the overall null distribution rather than any single permutation result, the estimate stabilizes quickly. Empirically, we observed that performing only one permutation produces a sufficiently stable null distribution and consistent thresholds, closely matching results from larger numbers of permutations. Therefore, SpaPheno sets one permutation as the default, while allowing users to increase this number if desired.

### Ablation analysis

A central design of SpaPheno is the use of cell type proportions as unified features for both bulk RNA-seq and spatial transcriptomics data, enabling biologically meaningful alignment across modalities. In addition, SpaPheno explicitly integrates spatial information through SPACE (spatially adjacent cell type embedding), which encodes local spatial context around each spot. These two components—cell type annotation and spatial information integration—are key to enhancing the interpretability and accuracy of phenotype prediction.

To further assess the contribution of each component, we performed ablation experiments using simulated datasets from osmFISH and STARmap, comparing the full SpaPheno pipeline against three alternative settings:

- Without cell type annotation: Instead of cell type proportions, both bulk and spatial transcriptomics were represented by raw gene expression features. For spatial data, we incorporated spatial information by computing the average gene expression across spatially adjacent spots (analogous to SPACE, but applied to gene expression instead of cell types).
- Without spatial information: Cell type proportions were used for both bulk and spatial transcriptomics, but no spatial embedding (i.e., no SPACE) was applied to the spatial data.
- Without spatial information and cell type annotation: Raw gene expression values were used as features for both bulk and spatial data, without any spatial integration or cell type deconvolution.

For each setting, we trained elastic net models to predict simulated phenotype scores and evaluated the performance using recall, precision, and F1-score, comparing predictions with ground-truth labels.

This ablation analysis demonstrates that both **c**ell type-based feature representation and spatial context embedding are essential for the superior performance of SpaPheno. Removing either component results in a notable drop in predictive accuracy, highlighting the necessity of integrating cellular and spatial dimensions in spatial phenotype modeling.

### Simulation data construction

To systematically evaluate the performance of SpaPheno, we conducted simulations based on single-cell spatial transcriptomics datasets (primarily osmFISH and STARmap), which include both cell-type annotations and anatomical layer labels.

Basically, there are three steps: (1) Binary phenotype simulation. We randomly selected two anatomical layers from each dataset to represent binary phenotypes. These layers exhibit distinct cell-type compositions and spatial structures, serving as biologically grounded surrogates for phenotype separation. (2) Pseudo-bulk construction using SPACE features. For each cell, we calculated SPACE features that capture local spatial context based on neighborhood cell-type composition. These SPACE profiles were then aggregated to simulate pseudo-bulk samples for each phenotype group. To increase biological complexity, a fraction of cells from other layers was also included to simulate background noise. (3) Simulating inter-sample heterogeneity. To model cohort-level variability, we applied mild perturbations to the SPACE matrices before aggregation. Specifically, each value was multiplied by a random factor uniformly sampled from [(1 – ε), (1 + ε)], with ε = 0.1 by default. Since the perturbation is multiplicative, non-negativity is preserved by construction. This process was repeated to generate 50 pseudo-bulk samples per phenotype group.

### Evaluation metrics

To quantitatively assess the predictive performance of SpaPheno in simulation studies, we employed three widely used classification metrics: precision, recall, and F1-score. These metrics were calculated by comparing the predicted phenotype-relevant cells with the known ground-truth labels derived from simulation design.

Specifically, in each simulation setting, cells within the selected phenotype layers were defined as true positives (TP). SpaPheno assigns a phenotype score to each cell, and cells with scores in the tail corresponding to the phenotype (default: top 0.1% for positively associated phenotypes, or bottom 0.1% for negatively associated phenotypes) were classified as predicted positives. Predicted positives that fall inside the selected phenotype layer were counted as true positives (TP); predicted positives outside the selected layer were counted as false positives (FP); cells inside the selected layer that were not classified as predicted positives were counted as false negatives (FN). Performance was quantified at the cell level using precision = TP / (TP + FP), recall = TP / (TP + FN), and F1-score = 2 × (precision × recall) / (precision + recall).

These metrics were averaged across all simulated phenotype classes and replicates. Evaluation was performed under both the full SpaPheno model and the ablation settings to enable a fair comparison of predictive accuracy under different model configurations.

Note that these quantitative metrics were only used in simulation-based evaluations, where the ground truth is explicitly defined. For real spatial transcriptomics datasets, model performance was assessed through case studies and biological interpretability rather than direct metric-based comparisons.

### Cell type enrichment score for stage- vs. survival-specific regions

In the analysis of primary liver cancer, we compared spatial domains associated with tumor stage (early vs. late) and survival risk (low-risk vs. high-risk). While most early-stage regions overlapped with low-risk-survival regions and late-stage regions overlapped with high-risk-survival regions, a small number of exceptions were observed (e.g., early-stage + high-risk-survival: 4 spots [0.4%]; late-stage + low-risk-survival: 23 spots [2.5%]). Due to their limited representation, these regions were excluded from downstream analysis.

We focused on stage-specific but survival-independent spatial regions and compared them to the corresponding stage-specific, risk-associated regions. To quantify the relative enrichment of cell types between these region pairs, we defined a Cell Type Enrichment Score (CTES).

Let 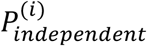 and 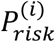 denote the proportion of a given cell type in the risk-independent and risk-associated regions, respectively, for spatial slice *i* ∈ {1, …, *n*}. The CTES is defined as:

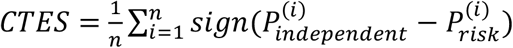

The CTES ranges from −1 to 1:

- CTES = 1: the cell type is more abundant in the risk-independent region in all slices;
- CTES = −1: it is more abundant in the risk-associated region across all slices;
- CTES = 0: no consistent enrichment pattern.

We applied this metric to:

- Early-stage-specific, risk-independent vs. early-stage-specific, low-risk-survival regions
- Late-stage-specific, risk-independent vs. late-stage-specific, high-risk-survival regions

This allowed us to identify cell types selectively enriched in tumor-stage-related spatial domains independent of survival phenotype.

### Data descriptions

We validated SpaPheno on two simulations on osmFISH and STARmap datasets and five cancer spatial transcriptomics (ST) with corresponding single-cell RNA-seq (scRNA-seq) reference, and bulk RNA-seq data with clinical phenotype labels. For all ST and bulk samples, cell2location was performed for deconvolution with default parameters. We chose cell2location due to its superior performance in benchmarks and comparative analyses in our previous study[6, 30]. Detailed information on the datasets used in this study is shown in **Supplementary Table 1** and summarized below:

#### Primary Liver Cancer

We used 6 ST slices with tertiary lymphoid structure (TLS) annotations was available at https://ngdc.cncb.ac.cn/gsa-human/browse/HRA000437. The scRNA-seq reference[16] was available at https://data.mendeley.com/datasets/skrx2fz79n/1. Matched bulk RNA-seq and clinical phenotypes (survival, stage) were derived from TCGA-LIHC: https://xenabrowser.net/datapages/?cohort=TCGA%20Liver%20Cancer%20(LIHC).

#### Hepatocellular Carcinoma (HCC)

We used 4 ST slices[16] and their matched scRNA-seq reference[16], with annotations for immune, fibroblast, and SPP1^+^ macrophage populations, available at https://data.mendeley.com/datasets/skrx2fz79n/1. Bulk RNA-seq and clinical data were obtained from TCGA-LIHC: https://xenabrowser.net/datapages/?cohort=TCGA%20Liver%20Cancer%20(LIHC).

#### Breast Cancer (BRCA)

We used 4 ST slices[18] with immune infiltration annotations, which was available at https://ega-archive.org/datasets/EGAD00001008031. The scRNA-seq reference[31] was available at https://www.ncbi.nlm.nih.gov/geo/query/acc.cgi?acc=GSE176078. Bulk RNA-seq and survival data were collected from TCGA-BRCA: https://xenabrowser.net/datapages/?cohort=TCGA%20Breast%20Cancer%20(BRCA).

#### Clear Cell Renal Cell Carcinoma (ccRCC)

We used 5 ST slices[17] with TLS annotations, which was available at https://www.ncbi.nlm.nih.gov/geo/query/acc.cgi?acc=GSE175540. The scRNA-seq reference[32] was available at https://data.mendeley.com/datasets/g67bkbnhhg/1. Bulk RNA-seq and clinical data were from TCGA-KIRC: https://xenabrowser.net/datapages/?cohort=TCGA%20Kidney%20Cancer%20(KIRC).

#### Melanoma

We used one ST slice with tumor and immune cell annotations, which was available at https://www.spatialresearch.org/resources-published-datasets/doi-10-1158-0008-5472-can-18-0747. The scRNA-seq reference[33] was available at https://www.ncbi.nlm.nih.gov/geo/query/acc.cgi?acc=GSE115978. Bulk RNA-seq and immune checkpoint blockade (ICB) response data[28] were available at https://www.ncbi.nlm.nih.gov/geo/query/acc.cgi?acc=GSE78220.

## Supporting information

Supplementary list

## Data availability

All data analyzed in this paper are available in raw form from their original studies.

## Code availability

The SpaPheno algorithm is implemented in R and is available on GitHub [https://github.com/Duan-Lab1/SpaPheno], along with detailed documentation, demo code, and data to facilitate reproducibility.

## Authors’ contributions

B.D. conceived the method. B.D., X.J.C., H.Z. implemented the pipeline and processed the data. B.D. wrote the manuscript with assistance from other authors.

## Fundings

This work was supported by the National Natural Science Foundation of China (Grant No. 61572361), Shanghai Rising-Star Program Sailing Special (Grant No. 23YF1450200), Shanghai Key Research and Development Program of Computational Biology (Grant No. 25JS2850100).

## Competing interests

The authors declare no competing interests.

## Notes

### Competing Interest Statement

The authors have declared no competing interest.

## References

1. Marx V: Method of the Year: spatially resolved transcriptomics. Nat Methods 2021, 18:9–14.

2. Moses L, Pachter L: Museum of spatial transcriptomics. Nat Methods 2022, 19:534–546.

3. Hao Y, Hao S, Andersen-Nissen E, Mauck WM, 3rd, Zheng S, Butler A, Lee MJ, Wilk AJ, Darby C, Zager M, et al: Integrated analysis of multimodal single-cell data. Cell 2021, 184:3573–3587 e3529.

4. Walker C, Angelo M: Toward clinical applications of spatial-omics in cancer research. Nat Cancer 2024, 5:1771–1773.

5. Regev A, Teichmann SA, Lander ES, Amit I, Benoist C, Birney E, Bodenmiller B, Campbell P, Carninci P, Clatworthy M, et al: The Human Cell Atlas. Elife 2017, 6.

6. Duan B, Chen S, Cheng X, Liu Q: Multi-slice spatial transcriptome domain analysis with SpaDo. Genome Biol 2024, 25:73.

7. Asp M, Giacomello S, Larsson L, Wu C, Furth D, Qian X, Wardell E, Custodio J, Reimegard J, Salmen F, et al: A Spatiotemporal Organ-Wide Gene Expression and Cell Atlas of the Developing Human Heart. Cell 2019, 179:1647–1660 e1619.

8. Velten B, Stegle O: Principles and challenges of modeling temporal and spatial omics data. Nat Methods 2023, 20:1462–1474.

9. Cheng X, Tang C, Dong K, You Y, Zhao X, Duan B, Chen S, Chuai G, Zhang Z, Liu Q: SpaLinker identifies phenotype-associated spatial tumor microenvironment features by integrating bulk and spatial sequencing data. Cell Genom 2025, 5:100893.

10. Zuo C, Xia J, Xu Y, Xu Y, Gao P, Zhang J, Wang Y, Chen L: stClinic dissects clinically relevant niches by integrating spatial multi-slice multi-omics data in dynamic graphs. Nat Commun 2025, 16:5317.

11. Zou H HT: Regularization and variable selection via the elastic net. Journal of the Royal Statistical Society Series B: Statistical Methodology 2005, 67:301–320.

12. Lundberg S M LSI: A unified approach to interpreting model predictions. Advances in neural information processing systems 2017:30.

13. Codeluppi S, Borm LE, Zeisel A, La Manno G, van Lunteren JA, Svensson CI, Linnarsson S: Spatial organization of the somatosensory cortex revealed by osmFISH. Nat Methods 2018, 15:932–935.

14. Wang X, Allen WE, Wright MA, Sylwestrak EL, Samusik N, Vesuna S, Evans K, Liu C, Ramakrishnan C, Liu J, et al: Three-dimensional intact-tissue sequencing of single-cell transcriptional states. Science 2018, 361.

15. Wu R, Guo W, Qiu X, Wang S, Sui C, Lian Q, Wu J, Shan Y, Yang Z, Yang S, et al: Comprehensive analysis of spatial architecture in primary liver cancer. Sci Adv 2021, 7:eabg3750.

16. Liu Y, Xun Z, Ma K, Liang S, Li X, Zhou S, Sun L, Liu Y, Du Y, Guo X, et al: Identification of a tumour immune barrier in the HCC microenvironment that determines the efficacy of immunotherapy. J Hepatol 2023, 78:770–782.

17. Meylan M, Petitprez F, Becht E, Bougouin A, Pupier G, Calvez A, Giglioli I, Verkarre V, Lacroix G, Verneau J, et al: Tertiary lymphoid structures generate and propagate anti-tumor antibody-producing plasma cells in renal cell cancer. Immunity 2022, 55:527–541 e525.

18. Andersson A, Larsson L, Stenbeck L, Salmen F, Ehinger A, Wu SZ, Al-Eryani G, Roden D, Swarbrick A, Borg A, et al: Spatial deconvolution of HER2-positive breast cancer delineates tumor-associated cell type interactions. Nat Commun 2021, 12:6012.

19. Thrane K, Eriksson H, Maaskola J, Hansson J, Lundeberg J: Spatially Resolved Transcriptomics Enables Dissection of Genetic Heterogeneity in Stage III Cutaneous Malignant Melanoma. Cancer Res 2018, 78:5970–5979.

20. Duan B, Zhu C, Chuai G, Tang C, Chen X, Chen S, Fu S, Li G, Liu Q: Learning for single-cell assignment. Sci Adv 2020, 6.

21. Kleshchevnikov V, Shmatko A, Dann E, Aivazidis A, King HW, Li T, Elmentaite R, Lomakin A, Kedlian V, Gayoso A, et al: Cell2location maps fine-grained cell types in spatial transcriptomics. Nat Biotechnol 2022, 40:661–671.

22. Sun D, Guan X, Moran AE, Wu LY, Qian DZ, Schedin P, Dai MS, Danilov AV, Alumkal JJ, Adey AC, et al: Identifying phenotype-associated subpopulations by integrating bulk and single-cell sequencing data. Nat Biotechnol 2022, 40:527–538.

23. Schumacher TN, Thommen DS: Tertiary lymphoid structures in cancer. Science 2022, 375:eabf9419.

24. Zhang Q, He Y, Luo N, Patel SJ, Han Y, Gao R, Modak M, Carotta S, Haslinger C, Kind D, et al: Landscape and Dynamics of Single Immune Cells in Hepatocellular Carcinoma. Cell 2019, 179:829–845 e820.

25. Zheng C, Zheng L, Yoo JK, Guo H, Zhang Y, Guo X, Kang B, Hu R, Huang JY, Zhang Q, et al: Landscape of Infiltrating T Cells in Liver Cancer Revealed by Single-Cell Sequencing. Cell 2017, 169:1342–1356 e1316.

26. Helmink BA, Reddy SM, Gao J, Zhang S, Basar R, Thakur R, Yizhak K, Sade-Feldman M, Blando J, Han G, et al: B cells and tertiary lymphoid structures promote immunotherapy response. Nature 2020, 577:549–555.

27. Paijens ST, Vledder A, de Bruyn M, Nijman HW: Tumor-infiltrating lymphocytes in the immunotherapy era. Cell Mol Immunol 2021, 18:842–859.

28. Hugo W, Zaretsky JM, Sun L, Song C, Moreno BH, Hu-Lieskovan S, Berent-Maoz B, Pang J, Chmielowski B, Cherry G, et al: Genomic and Transcriptomic Features of Response to Anti-PD-1 Therapy in Metastatic Melanoma. Cell 2016, 165:35–44.

29. Zhao E, Stone MR, Ren X, Guenthoer J, Smythe KS, Pulliam T, Williams SR, Uytingco CR, Taylor SEB, Nghiem P, et al: Spatial transcriptomics at subspot resolution with BayesSpace. Nat Biotechnol 2021, 39:1375–1384.

30. Li B, Zhang W, Guo C, Xu H, Li L, Fang M, Hu Y, Zhang X, Yao X, Tang M, et al: Benchmarking spatial and single-cell transcriptomics integration methods for transcript distribution prediction and cell type deconvolution. Nat Methods 2022, 19:662–670.

31. Wu SZ, Al-Eryani G, Roden DL, Junankar S, Harvey K, Andersson A, Thennavan A, Wang C, Torpy JR, Bartonicek N, et al: A single-cell and spatially resolved atlas of human breast cancers. Nat Genet 2021, 53:1334–1347.

32. Bi K, He MX, Bakouny Z, Kanodia A, Napolitano S, Wu J, Grimaldi G, Braun DA, Cuoco MS, Mayorga A, et al: Tumor and immune reprogramming during immunotherapy in advanced renal cell carcinoma. Cancer Cell 2021, 39:649–661 e645.

33. Jerby-Arnon L, Shah P, Cuoco M, Rodman C, Su M, Melms J, Leeson R, Kanodia A, Mei S, Lin J: Single-cell RNA-seq of melanoma ecosystems reveals sources of T cells exclusion linked to immunotherapy clinical outcomes. Gene Expr Omnibus 2018.

