## Supplementary list for "SpaPheno: Linking Spatial Transcriptomics to Clinical Phenotypes with Interpretable Machine Learning"

### Supplementary Materials for "SpaPheno: Linking Spatial Transcriptomics to Clinical Phenotypes with Interpretable Machine Learning".

#### This file includes:

#### Supplementary Table 1

#### Supplementary Fig. 1-11

**Supplementary Table 1: Datasets Used in This Study**

| Cancer Type | Data Type | Description | Reference | Data Access Link |
|---|---|---|---|---|
| **ccRCC** | ST dataset | Spatial transcriptomics of clear cell renal cell carcinoma | Meylan et al., *Immunity* (2022) | https://www.ncbi.nlm.nih.gov/geo/query/acc.cgi?acc=GSE175540 |
| **ccRCC** | scRNA-seq reference | Non-ICB-treated samples (P76 and P90) from advanced ccRCC | Bi et al., *Cancer Cell* (2021) | https://data.mendeley.com/datasets/g67bkbnhhg/1 |
| **Primary liver cancer** | ST dataset | Spatial architecture of primary liver cancer | Wu et al., *Science Advances* (2021) | https://ngdc.cncb.ac.cn/gsa-human/browse/HRA000437 |
| **Primary liver cancer** | scRNA-seq reference | Tumor immune microenvironment of HCC | Liu et al., *Journal of Hepatology* (2023) | https://data.mendeley.com/datasets/skrx2fz79n/1 |
| **HCC** | ST dataset | HCC spatial transcriptomics (from same Liu et al. 2023 study) | Liu et al., *Journal of Hepatology* (2023) | https://data.mendeley.com/datasets/skrx2fz79n/1 |
| **HCC** | scRNA-seq | Single-cell reference of | Liu et al., *Journal of* | https://data. |

| Cancer Type | Data Type | Description | Reference | Data Access Link |
|---|---|---|---|---|
| | reference | HCC immune landscape | *Hepatology* (2023) | mendeley.com/datasets/skrx2fz79n/1 |
| **BRCA** | ST dataset | HER2-positive breast cancer spatial transcriptomics | Andersson et al., *Nature Communications* (2021) | https://ega-archive.org/datasets/EGAD00001008031 |
| **BRCA** | scRNA-seq reference | Single-cell and spatial atlas of breast cancer | Wu et al., *Nature Genetics* (2021) | https://www.ncbi.nlm.nih.gov/geo/query/acc.cgi?acc=GSE176078 |
| **Melanoma** | ST dataset | Stage III melanoma spatial transcriptomics | Thrane et al., *Cancer Research* (2018) | https://www.spatialresearch.org/resources-published-datasets/doi-10-1158-0008-5472-can-18-0747/ |
| **Melanoma** | scRNA-seq reference | Melanoma immune ecosystem and ICB response | Jerby-Arnon et al., *GEO* (2018) | https://www.ncbi.nlm.nih.gov/geo/query/acc.cgi?acc=GSE115978 |
| **KIRC (TCGA)** | Bulk RNA-seq | Kidney renal clear cell carcinoma | UCSC Xena | https://xena.ucsc.edu/ |
| **LIHC (TCGA)** | Bulk RNA-seq | Liver hepatocellular carcinoma | UCSC Xena | https://xena.ucsc.edu/ |
| **BRCA (TCGA)** | Bulk RNA-seq | Breast invasive carcinoma | UCSC Xena | https://xena.ucsc.edu/ |
| **Melanoma** | Bulk RNA-seq (ICB) | Response to anti–PD-1 therapy in metastatic melanoma | Hugo et al., *Cell* (2016) | https://www.ncbi.nlm.nih.gov/geo/query/acc.cgi?acc=GSE78220 |

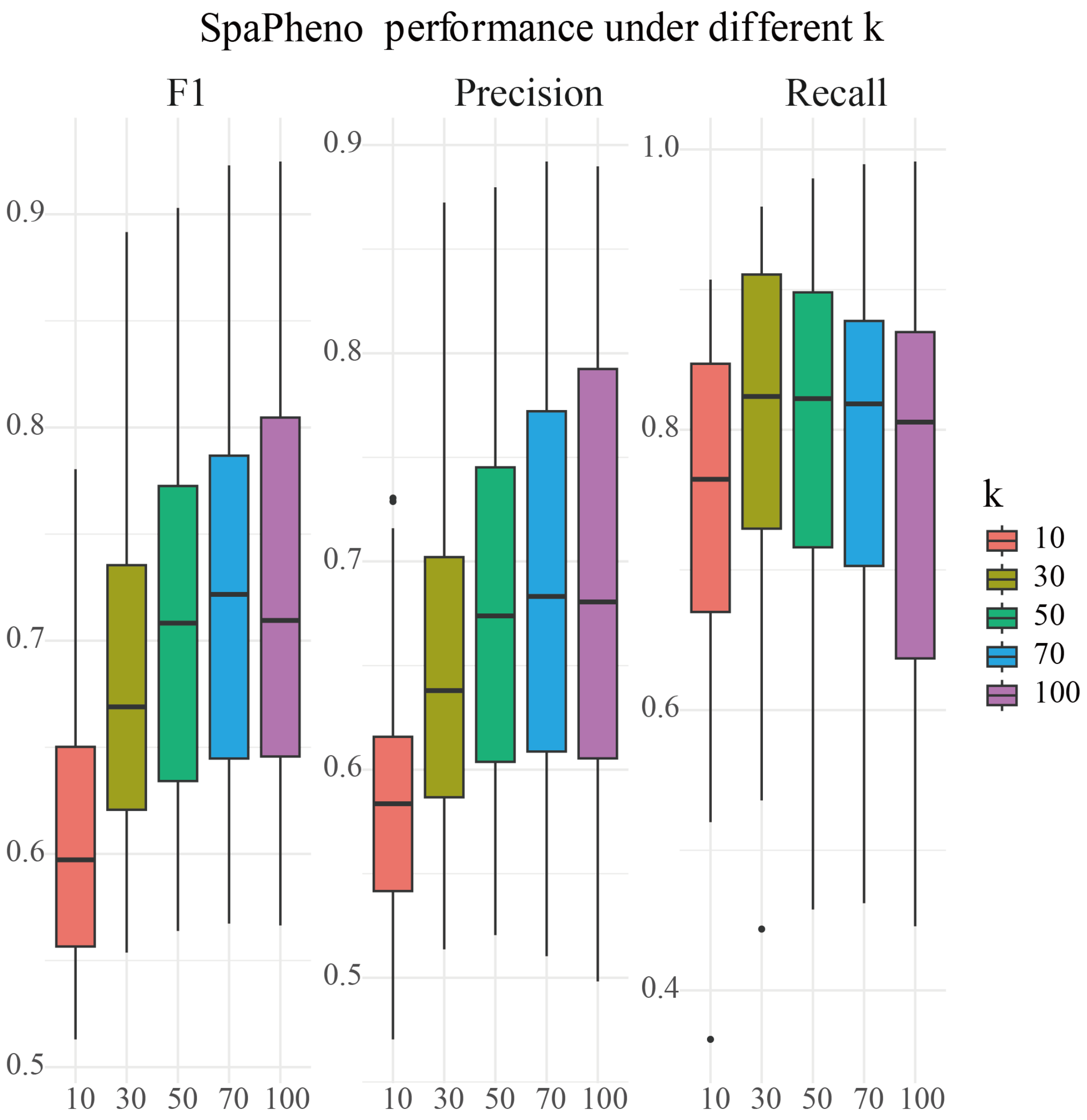


**Supplementary Fig. 1 | SpaPheno performance under different k for choosing neighbors on osmFISH data.**

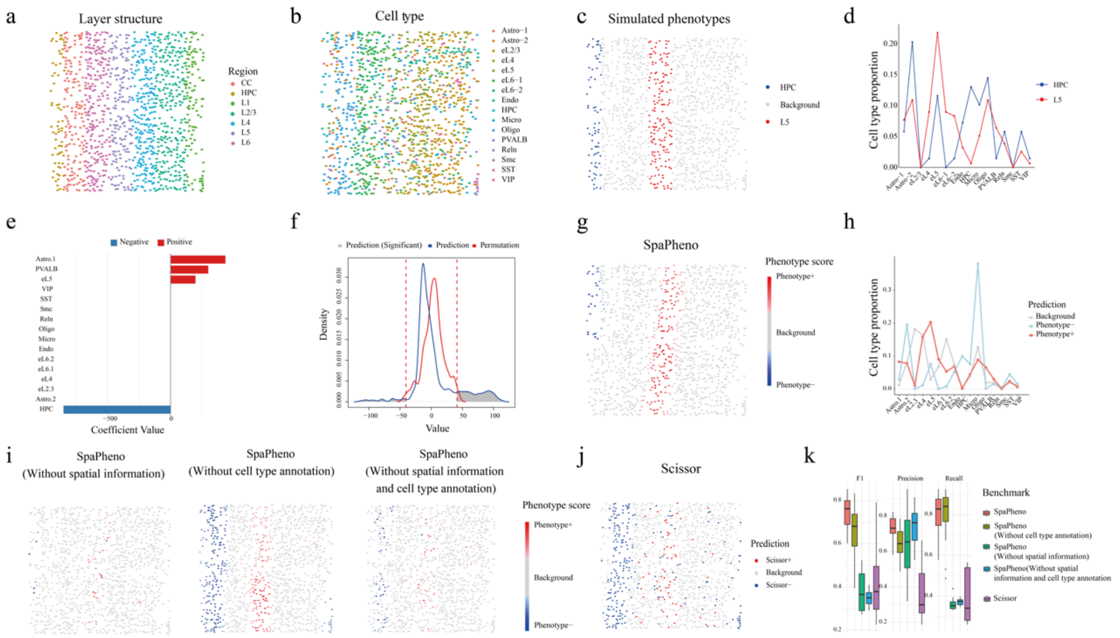


**Supplementary Fig. 2 | Evaluation of SpaPheno on simulated phenotypes using real STARmap data.** **a–b**, Layer structure and cell type annotations in the STARmap dataset. **c**, Simulated phenotype labels derived from STARmap data. **d**, Cell type composition of simulated phenotype groups. **e**, Global feature attributions based on model coefficients from SpaPheno. **f,** Distribution of SpaPheno-predicted phenotype scores compared with permutation controls. **g**, Spatial prediction map of phenotype scores across the tissue. **h**, Cell type composition of SpaPheno-predicted phenotype groups. **i,** Performance of SpaPheno under ablation settings by removing spatial information, cell-type information, or both. **j,** Prediction results from Scissor. **k,** Overall performance comparison of SpaPheno, its ablation variants, and Scissor across all simulated phenotype scenarios.

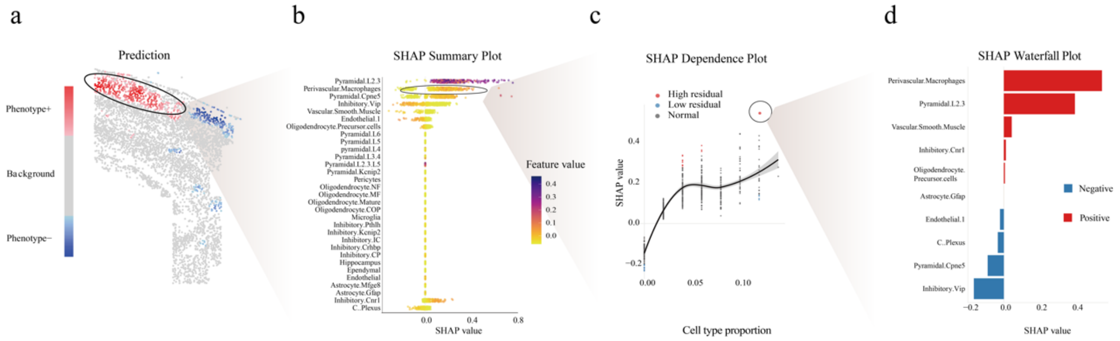


**Supplementary Fig. 3 | Multi-scale interpretability of SpaPheno via SHAP. a**, Spatial prediction map of phenotype scores across the tissue, showing the distribution of predicted phenotype. **b**, SHAP summary plot for phenotype+ spots. Each dot represents a spot, colored by the relative abundance of the corresponding cell type; positive SHAP values indicate features that contribute to classifying a spot as phenotype+, while negative values indicate the opposite. In this example, the phenotype corresponds to enrichment in the Layer3-median region. **c**, SHAP dependence plot for the cell type "Perivascular Macrophages," showing how its SHAP value changes with its proportion across spots. This highlights both high-residual (outlier) and low-residual (conserved) spots. **d**,

SHAP waterfall plot for a representative high-residual spot, illustrating the contribution (direction and magnitude) of each cell type to the final prediction.

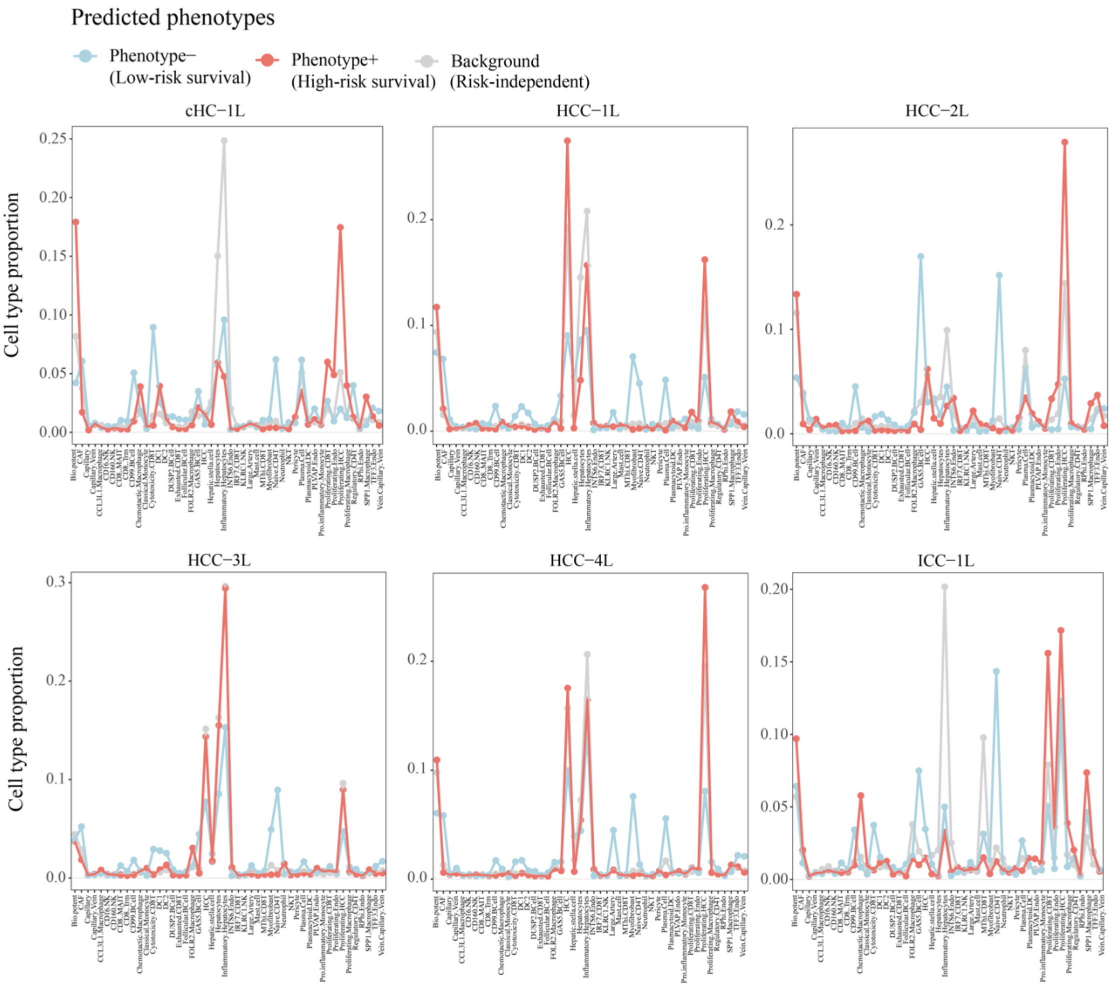


**Supplementary Fig. 4 | Cell type composition of SpaPheno-predicted risk survival associated regions across six primary liver cancer slices.**

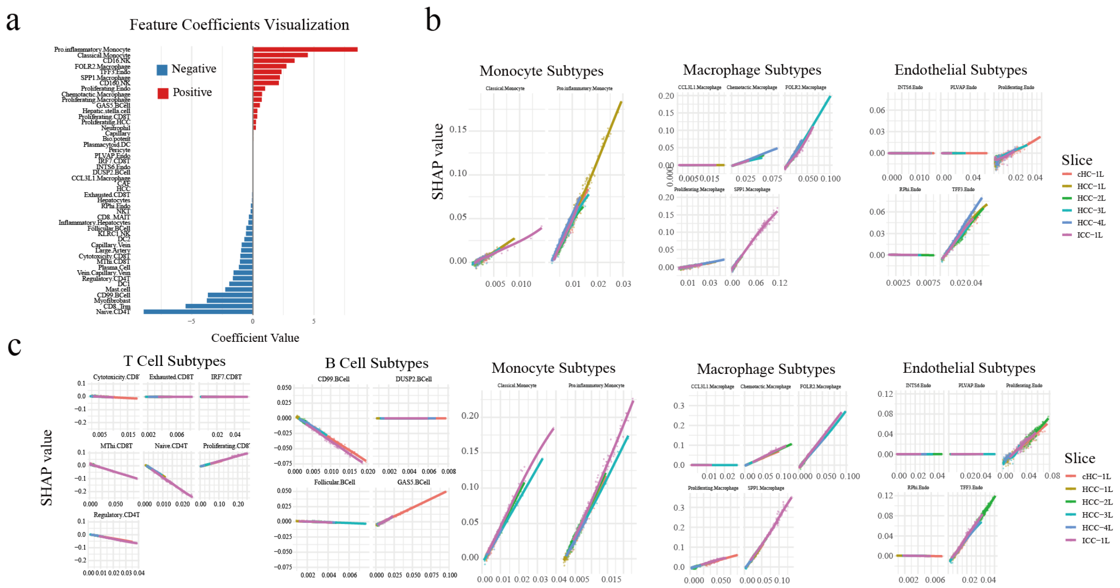


**Supplementary Fig. 5 | Global feature attributions and SHAP dependence plots for cell type subtypes associated with risk survival across six primary liver cancer slices. a**, Global feature attributions based on model coefficients derived from SpaPheno, highlighting the most predictive cell type subtypes across all slices. **b**, SHAP dependence plots for monocyte subtypes, macrophage subtypes, and endothelial subtypes associated with the low-risk survival phenotype. **c**, SHAP dependence plots for T cell subtypes, B cell subtypes, monocyte subtypes, macrophage subtypes, and endothelial subtypes associated with the high-risk survival phenotype. These plots illustrate how SHAP values vary with cell type proportions and reveal subtype-specific contributions to phenotype prediction.

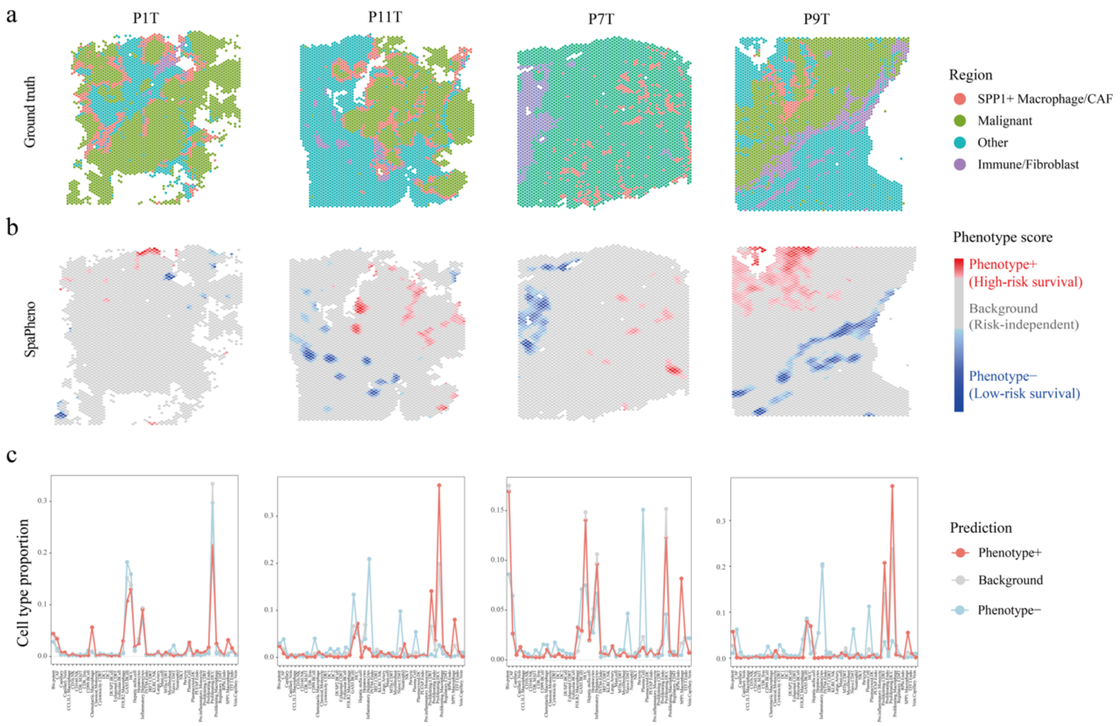


**Supplementary Fig. 6 | SpaPheno identifies immune-enriched regions associated with low-risk survival and SPP1$^{+}$ macrophage–enriched regions associated with high-risk survival in an independent HCC dataset. a–b**, Region annotations and SpaPheno-predicted survival-associated regions across 4 spatial transcriptomics (ST) slices. **c**, Cell type composition of SpaPheno-predicted phenotype regions across the 4 HCC slices.

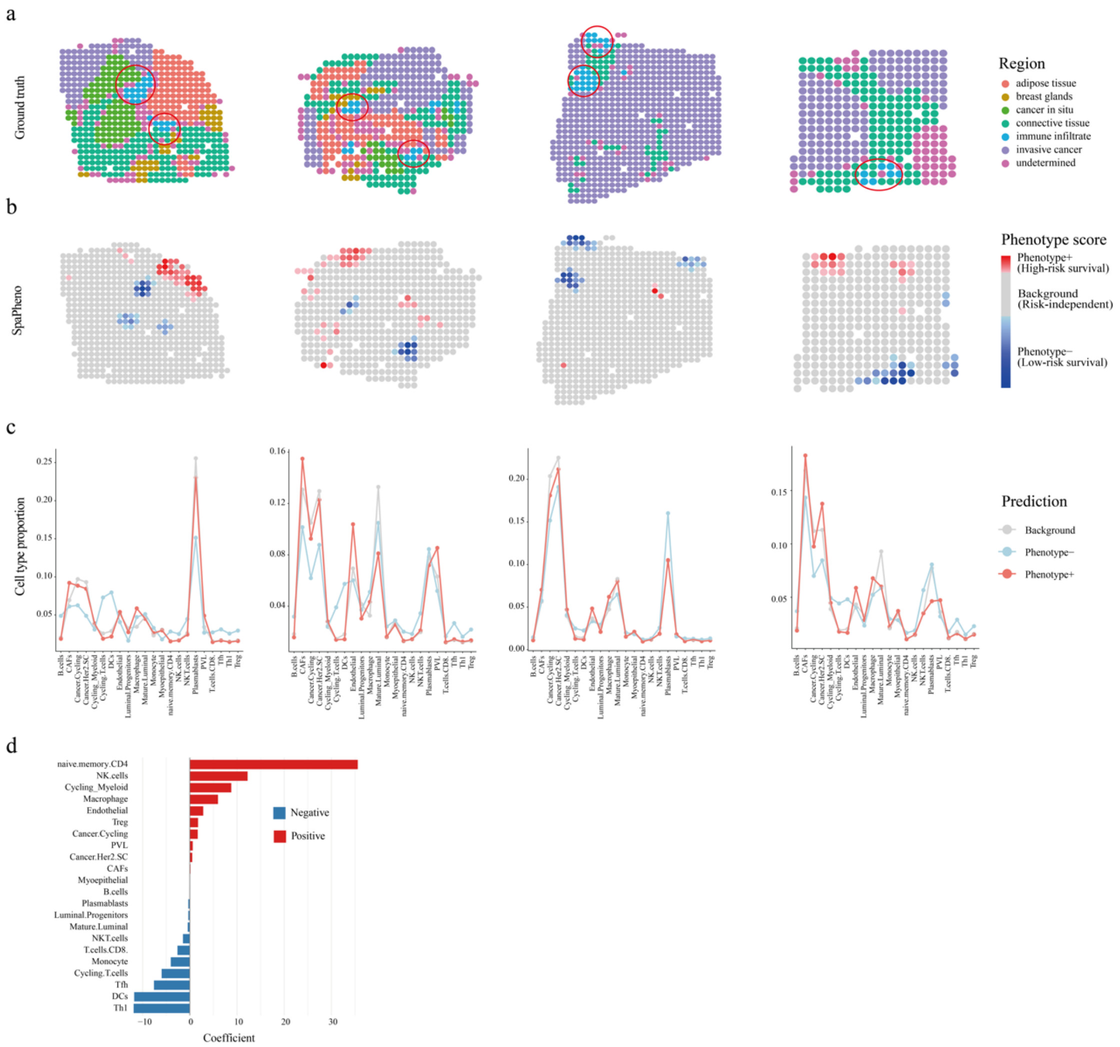


**Supplementary Fig. 7 | SpaPheno identifies immune-enriched regions associated with low-risk survival in a BRCA dataset. a–b**, Region annotations and SpaPheno-predicted survival-associated regions across 4 spatial transcriptomics (ST) slices. **c**, Cell type composition of SpaPheno-predicted phenotype regions across the 4 BRCA slices. **d**, Global feature attributions based on model coefficients derived from SpaPheno, highlighting the most predictive cell type subtypes across all slices.

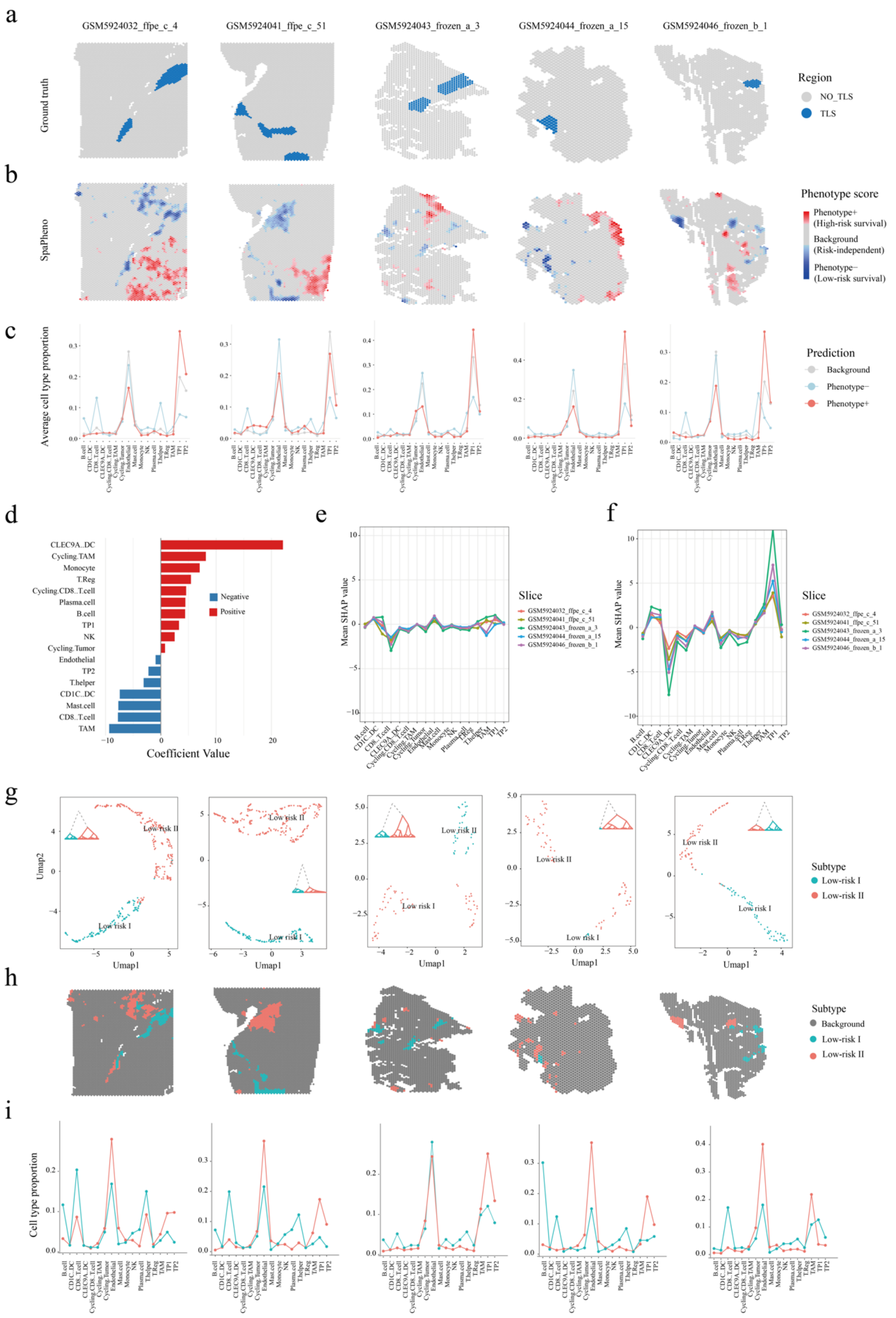


**Supplementary Fig. 8 | SpaPheno identifies TLS-like regions and endothelial-rich regions associated with low-risk survival in ccRCC. a–b**, Region annotations and SpaPheno-predicted survival-associated regions across 5 spatial transcriptomics (ST) slices. **c**, Cell type composition of SpaPheno-predicted phenotype regions across the 5 ccRCC ST slices. **d**, Global feature attributions based on model coefficients derived from SpaPheno, highlighting the most predictive cell type subtypes across all slices. **e**, Mean SHAP values of each cell type in predicted low-risk survival regions. **f**, Mean SHAP values of each cell type in predicted high-risk survival regions. **g**, UMAP visualization of two subtypes of low-risk survival regions. **h**, The two subtypes of SpaPheno-predicted low-risk survival-associated regions across 5 spatial transcriptomics (ST) slices. **i,** Cell type composition of SpaPheno-predicted phenotype regions across the 5 ccRCC slices.

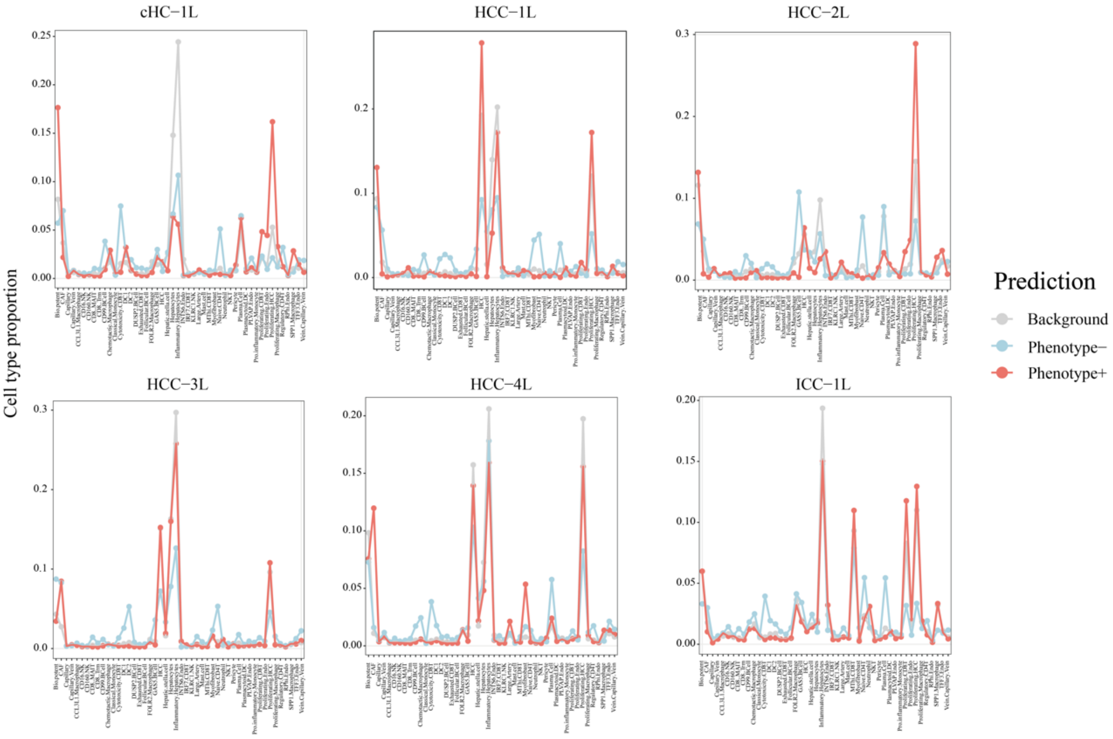


**Supplementary Fig. 9 | Cell type composition of SpaPheno-predicted tumor-stage associated regions across six primary liver cancer slices.**

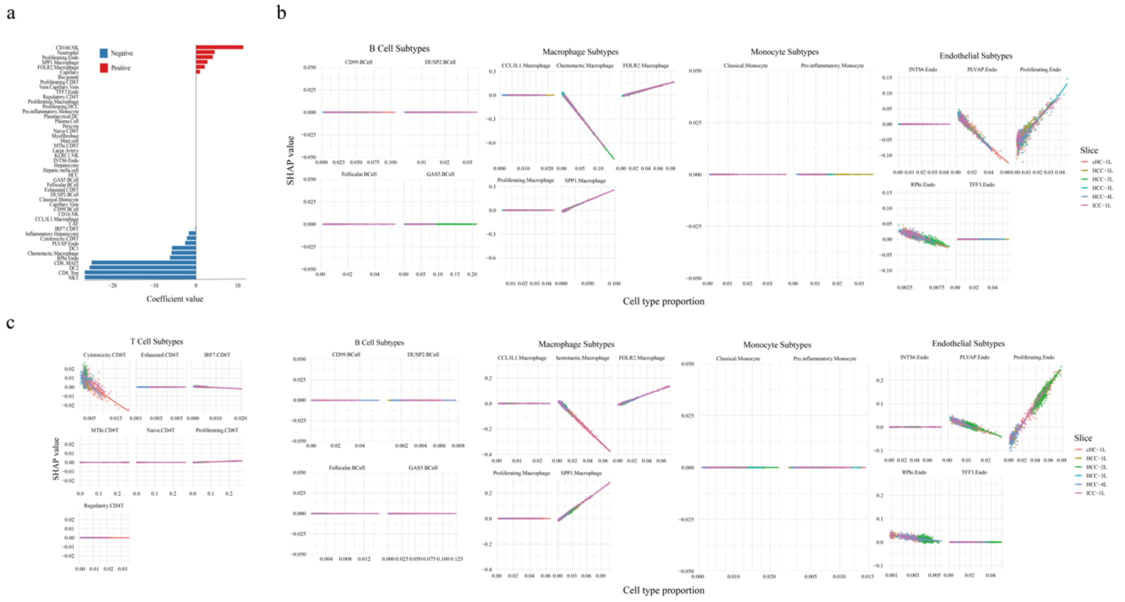

**Supplementary Fig. 10 | Global feature attributions and SHAP dependence plots for cell type subtypes associated with tumor stage across six primary liver cancer slices. a**, Global feature attributions based on model coefficients derived from SpaPheno, highlighting the most predictive cell type subtypes across all slices. **b**, SHAP dependence plots for B cell subtypes,monocyte subtypes, macrophage subtypes, and endothelial subtypes associated with the early-stage phenotype. **c**, SHAP dependence plots for T cell subtypes, B cell subtypes, monocyte subtypes, macrophage subtypes, and endothelial subtypes associated with the late-stage phenotype. These plots illustrate how SHAP values vary with cell type proportions and reveal subtype-specific contributions to phenotype prediction.

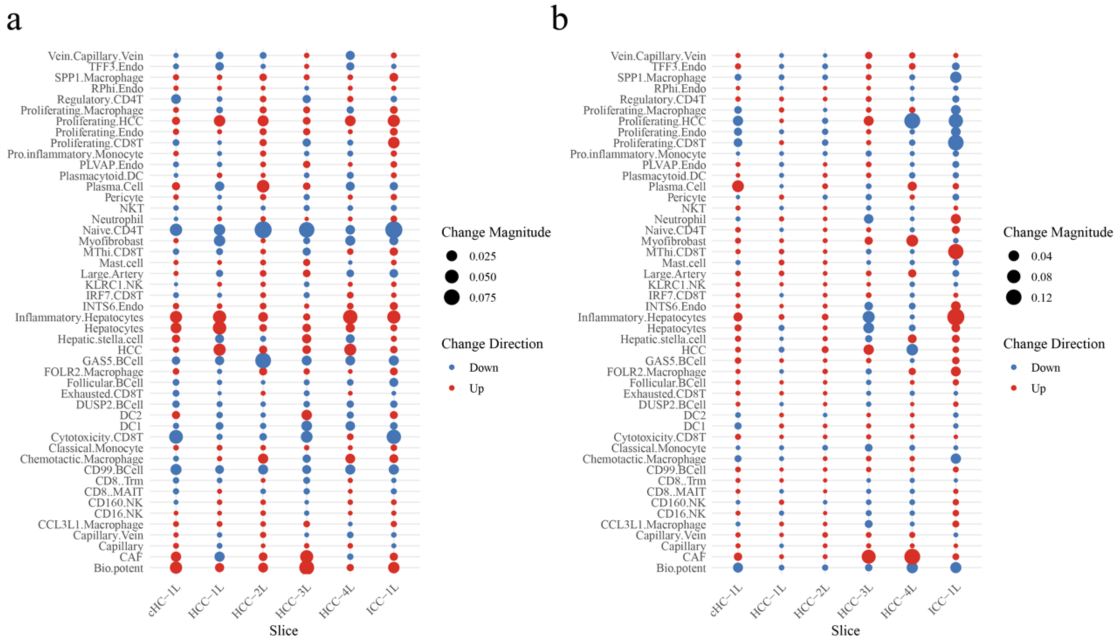


**Supplementary Fig. 11 | Cell type enrichment analysis of stage-specific but survival-independent regions. a**, Cell type enrichment of early-stage-specific, survival-independent regions compared with early-stage, low-risk survival-associated regions. **b**, Cell type enrichment of late-stage-specific, survival-independent regions compared with late-stage, high-risk survival-associated regions.
